# The rocaglate CR-31-B (-) inhibits SARS-CoV-2 replication at non-cytotoxic, low nanomolar concentrations *in vitro* and *ex vivo*

**DOI:** 10.1101/2020.11.24.389627

**Authors:** Christin Müller, Wiebke Obermann, Nadja Karl, Hans-Guido Wendel, Gaspar Taroncher-Oldenburg, Stephan Pleschka, Roland K. Hartmann, Arnold Grünweller, John Ziebuhr

## Abstract

Severe acute respiratory syndrome coronavirus 2 (SARS-CoV-2), a betacoronavirus in the subgenus *Sarbecovirus* causes a respiratory disease with varying symptoms referred to as coronavirus disease 2019 (COVID-19) and is responsible for a pandemic that started in early 2020. With no vaccines or effective antiviral treatments available, and infection and fatality numbers continuing to increase globally, the quest for novel therapeutic solutions remains an urgent priority. Rocaglates, a class of plant-derived cyclopenta[*b*]benzofurans, exhibit broad-spectrum antiviral activity against positive- and negative-sense RNA viruses. This compound class inhibits eukaryotic initiation factor 4A (eIF4A)-dependent mRNA translation initiation, resulting in strongly reduced viral RNA translation. The synthetic rocaglate CR-31-B (-) has previously been shown to inhibit the replication of human coronaviruses, such as HCoV-229E and MERS-CoV, as well as Zika-, Lassa-, Crimean Congo hemorrhagic fever virus in primary cells. Here, we assessed the antiviral activity of CR-31-B (-) against SARS-CoV-2 using both *in vitro* and *ex vivo* cell culture models. In African green monkey Vero E6 cells, CR-31-B (-) inhibited SARS-CoV-2 replication with an EC_50_ of ~1.8 nM. In line with this, viral protein accumulation and replication/transcription complex formation were found to be strongly reduced by this compound. In an *ex vivo* infection system using human airway epithelial cells, CR-31-B (-) was found to cause a massive reduction of SARS-CoV-2 titers by about 4 logs to nearly non-detectable levels. The data reveal a potent anti-SARS-CoV-2 activity by CR-31-B (-), corroborating previous results obtained for other coronaviruses and supporting the idea that rocaglates may be used in first-line antiviral intervention strategies against novel and emerging RNA virus outbreaks.

## 1. Introduction

Severe acute respiratory syndrome coronavirus 2 (SARS-CoV-2) is a positive-sense, single-stranded RNA betacoronavirus of zoonotic origin first detected in Wuhan, China, at the end of 2019. The virus causes a potentially severe respiratory illness, coronavirus disease 2019 (COVID-19), with infection fatality rates ranging from 0.5% to 1% according to recent estimates (Rajgor et al., 2020). Following its rapid spread first to Europe and then to the rest of the world, the World Health Organization declared SARS-CoV-2 a pandemic in March 2020 (WHO, 2020). SARS-CoV-2 is genetically very closely related to SARS-CoV, the causative agent of the 2002-2003 SARS outbreak (Wu et al., 2020, Zhou et al., 2020, Gorbalenya et al., 2020), and the two viruses have been assigned to the same virus species *Severe acute respiratory syndrome-related coronavirus* (Gorbalenya et al., 2020). The ongoing SARS-CoV-2 outbreak is the third documented case of a coronavirus outbreak of zoonotic origin to reach epidemic or pandemic scale since the beginning of the century (Gorbalenya et al., 2020). The SARS-CoV outbreak in 2002-2003 was followed by the Middle East respiratory syndrome coronavirus (MERS-CoV) outbreak in 2012/2013 (which is still ongoing) and the current SARS-CoV-2 outbreak, all of which can lead to severe pneumonia and death in humans (Anderson and Baric, 2012) and have caused substantial social and economic disruptions. To date, no approved therapeutic is available against SARS-CoV, MERS-CoV or SARS-CoV-2, but a number of investigational antiviral compounds that target viral functions, e.g. remdesivir, ribavirin and favipiravir, which target viral RNA synthesis, as well as several protease inhibitors have entered clinical trials (Brown et al., 2019, de Wit et al., 2020, Sheahan et al., 2017, de Wilde et al., 2014, Choy et al., 2020, Sanders et al., 2020). Moreover, compounds that modulate the human immune system and/or have an anti-inflammatory effect, including tocilizumab, interferon-beta and dexamethasone are also being tested for their potential to reduce the severity of COVID-19 progression (Oldenburg and Dohan, 2020, Guaraldi et al., 2020, Jalkanen et al., 2020, Lam et al., 2020). Finally, antivirals that target host mechanisms critical to viral replication and production of infectious virus progeny are also being developed, including inhibitors targeting host proteases required to activate the fusogenic activity of the SARS-CoV-2 spike protein (Bestle et al., 2020, Hoffmann et al., 2020).

Several groups, including ours, have focused on suppressing viral protein synthesis, another host function critical to viral proliferation. Incidentally, a recently published SARS-CoV-2 protein interaction map identified the host translational machinery as a top target for repurposing drugs to block SARS-CoV-2 (Gordon et al., 2020). In particular, two eukaryotic factors involved in the initiation phase of translation, eukaryotic initiation factor-1A (eIF1A) and eukaryotic initiation factor-4A (eIF4A), have been singled out for further development. eIF1A is a small protein that binds to the 40S ribosome subunit-mRNA complex (Gordon et al., 2020), and eIF4A is a DEAD-box RNA helicase central to the activity of the eukaryotic translation initiation complex eIF4F (Chu and Pelletier, 2015). As of this writing, the eIF1A inhibitor aplidin (PharmaMar) has entered a Phase 2 clinical trial for the treatment of COVID-19, and the eIF4A inhibitor zotatifin (eFFECTOR Therapeutics) is undergoing evaluation for potential COVID-19 clinical development (Harrison, 2020, Bronstrup and Sasse, 2020).

We have recently shown the potential of rocaglates, a class of natural and synthetic compounds characterized by a common cyclopenta[*b*]benzofuran skeleton and originally extracted from plants in the genus *Aglaia* (*Meliaceae*) (Ebada et al., 2011), to inhibit both the common cold human coronavirus HCoV-229E and MERS-CoV *in vitro* and *ex vivo* (Müller et al., 2018a, Müller et al., 2020). Natural rocaglates such as rocaglamide A (RocA) or silvestrol, and synthetic rocaglates such as CR-31-B (-) and the above-mentioned zotatifin are specific nanomolar eIF4A inhibitors. Rocaglates form stacking interactions with polypurine sequences in the 5′-untranslated regions (UTRs) of capped mRNAs, clamping the mRNAs onto eIF4A and stalling mRNA unwinding, which results in depletion of the mRNA-eIF4A complex from eIF4F. Because eIF4A-mediated translation is key to the activation of a majority of oncogenes, several rocaglates are in advanced preclinical cancer studies and at least one compound, zotatifin, has been advanced into early stage clinical studies (Bordeleau et al., 2008, Lucas et al., 2009, Kogure et al., 2013, Patton et al., 2015, Ernst et al., 2020).

Many viral RNAs contain structured 5’-UTRs analogous to eukaryotic capped mRNA UTRs and are dependent on eIF4A for translation (Madhugiri et al., 2016, Schlereth et al., 2016). Silvestrol exhibits broad-spectrum antiviral activity against a range of positive- and negative-sense RNA viruses such as Ebola virus, Zika virus, Chikungunya virus, Crimean Congo hemorrhagic fever virus, Lassa virus, hepatitis E and several coronaviruses (Biedenkopf et al., 2017, Elgner et al., 2018, Henss et al., 2018, Müller et al., 2020, Glitscher et al., 2018, Todt et al., 2018, Müller et al., 2018a). While silvestrol and other natural rocaglates represent promising therapeutic leads due to their low nanomolar activities and high selectivity indices (≥ 100), their natural availability is limited (Pannell, 1998), and their chemical synthesis is not optimal due to their complex structures (Adams et al., 2009, Pan et al., 2014). As an alternative, a multiplicity of synthetic rocaglate analogs have been generated that exhibit similar or enhanced eIF4A-targeting characteristics and can be produced to high purity under GLP conditions and in large quantities.

Recently, we compared the antiviral activity of silvestrol with that of CR-31-B (-), which lacks the dioxane moiety of silvestrol and makes it structurally less complex and much more straightforward to synthesize (Wolfe et al., 2014). We showed that CR-31-B (-) exhibits levels of viral replication inhibition analogous to those of silvestrol across a range of RNA viruses both *in vitro* and *ex vivo* (Müller et al., 2020). So far, zotatifin has only been tested on one coronavirus, SARS-CoV-2, and only in the Vero E6 cell line, showing similar efficacy as silvestrol and CR-31-B (-) (Gordon et al., 2020).

Here, we evaluated the potential activity of the synthetic rocaglate CR-31-B (-) against SARS-CoV-2 using both *in vitro* and human *ex vivo* cell culture systems.

## 2. Materials and methods

### 2.1 Cell culture

Vero E6 cells were grown in Dulbecco’s modified Eagle’s medium (DMEM) supplemented with 10% fetal bovine serum (FBS), 100 U/ml penicillin, and 100 μg/ml streptomycin at 37 °C in an atmosphere containing 5% CO_2_. HepG2 cells were cultured in Iscove’s Modified Dulbecco’s Medium (IMDM) supplemented with 10 % fetal calf serum (FCS) at 37 °C and 5 % CO_2_.

### 2.2 Reagents

Silvestrol was obtained from the Sarawak Biodiversity Centre (Kuching; North-Borneo, Malaysia; purity > 99%). A 6 mM stock solution was prepared in DMSO (sterile-filtered; Roth) and diluted in DMEM or IMDM. Control cells were treated with correspondingly concentrated DMSO dilutions lacking silvestrol. CR-31-B (-) and CR-31-B (+) (Wolfe et al., 2014) were dissolved in DMSO at a concentration of 10 mM and stored at −20 °C.

### 2.3 Dual luciferase reporter assay

The dual luciferase assay was performed as described previously in at least three independent replicates (Müller et al., 2018a, Müller et al., 2020). The total length of the analyzed 5’-UTRs, including single-stranded and double-stranded regions, ranges from 50 bp to 292 bp (ß-globin: 50 bp, (AC)_15_: 30 bp, (AG)_15_: 30 bp, HCoV-229E: 292 bp, MERS-CoV: 278 bp, SARS-CoV-2: 265 bp).

### 2.4 Cell toxicity

Cell viability of Vero E6 cells in the presence of the respective compounds was determined by MTT assay as described previously (Müller et al., 2018a).

### 2.5 Human airway epithelial cells

Cryopreserved normal human bronchial epithelial (NHBE) cells were obtained from Lonza. The undifferentiated cells were seeded on collagen IV-coated transwell plates (Corning Costar) and grown in a mixture of DMEM (Invitrogen) and BEGM (Lonza) supplemented with retinoic acid (75 nM). Every other day, fresh medium was added and, after reaching confluence, the cells were cultivated under air-liquid conditions for at least four additional weeks to differentiate into a pseudostratified human airway epithelium. During this period, medium from the basolateral compartment was renewed every 2–3 days and the apical surface was washed once a week with PBS (Invitrogen).

### 2.6 Antiviral activity

To determine the 50% effective concentration (EC_50_) for CR-31-B (-), Vero E6 cells were inoculated with SARS-CoV-2 (kindly provided by Christian Drosten) at a multiplicity of infection (MOI) of 0.1 at 33 °C. After 1 h, the inoculum was removed and cells were incubated with fresh medium containing CR-31-B (-) at increasing concentrations. Virus-containing supernatants were collected at 24 h post infection (p.i.) and virus titers were analyzed via plaque assay. EC_50_ values were determined based on virus titers in supernatants of infected cells treated with solvent control (DMSO) compared to virus titers in supernatants of infected cells treated with the respective inhibitor concentration. EC_50_ values were then calculated by non-linear regression analysis using GraphPad Prism 6.0 (GraphPad Software).

For the infection of NHBE cells, the apical surface was washed 3 times with PBS and cells were infected with SARS-CoV-2 (MOI = 3). After 1 h, the inoculum was removed and the medium in the basal compartment was replaced with medium containing the indicated inhibitor concentration. At the indicated time points p.i., the apical surface of the cells was incubated with PBS for 15 min and virus titers in the supernatants were determined by plaque assay.

### 2.7 Western blot analysis

To analyze viral protein accumulation, Vero E6 cells were infected with SARS-CoV-2 at an MOI of 1. After inoculation, the supernatant was replaced with fresh medium supplemented with the indicated concentrations of the respective CR-31-B enantiomer. After 24 h, the medium was removed, the cells were washed with PBS and lysed using buffer containing 50 mM Tris-HCl, pH 7.5, 150 mM NaCl, 1% NP40, and 1x protease inhibitor cocktail (P8340; Sigma-Aldrich). The insoluble material was removed by centrifugation and the protein content in the supernatant was measured using a Qubit 3 fluorometer (Invitrogen) and equal amounts of proteins were separated in SDS-10 % polyacrylamide gels and blotted onto a nitrocellulose membrane (Amersham). Membranes were incubated with polyclonal rabbit anti-SARS nucleocapsid protein antibody (Rockland) and mouse-anti actin antibody (abcam), respectively, each diluted 1:500 in PBS containing 1 % bovine serum albumin (BSA). After 60 min, membranes were washed with PBS and incubated with appropriate secondary antibodies (IRDye-conjugated anti-mouse or anti-rabbit IgG mAb [Li-COR Biosciences]) diluted 1:10.000 in PBS containing 1% BSA. After 1 h, membranes were washed and analyzed using the LI-COR Odyssey imaging system.

### 2.8 Immunofluorescence

Immunofluorescence was performed as described previously (Müller et al., 2018b). Briefly, Vero E6 cells were infected with SARS-CoV-2 (MOI of 1) and treated with the indicated concentrations of CR-31-B (-) or CR-31-B (+) for 24 hpi or left untreated. Then, the cells were fixed with ice-cold methanol and stained with mouse anti-dsRNA mAb (J2, SCICONS English & Scientific Consulting Kft.). As secondary antibodies, AlexaFluor 594 goat anti-mouse IgG was used. Confocal microscopy was done using a Leica SP05 CLSM and LAS-AF software (Leica).

## 3. Results

### 3.1. Inhibitory effect of CR-31-B (-) on eIF4A-dependent translation of viral 5’-UTRs

In a first set of experiments, we analyzed the inhibitory effect of CR-31-B (-) (Fig. 1A) on 5’-UTRs of different coronaviruses including SARS-CoV-2 in a dual luciferase reporter assay (Fig. 1B) to assess whether translation of mRNAs containing these viral 5’-UTRs depends on eIF4A. Beta-globin, as well as an unstructured (AC)_15_ sequence served as negative controls, whereas the polypurine sequence (AG)_15_ was used as a positive control since this sequence can be efficiently clamped by different rocaglates due to π-π stacking interactions on the surface of eIF4A-RNA complexes (Iwasaki et al., 2019, Müller et al., 2020). The data showed that the 5’-UTRs of SARS-CoV-2, HCoV-229E and MERS-CoV were similarly sensitive to translation inhibition by CR-31-B (-) in the dual luciferase reporter assay, when 10 nM CR-31-B (-) was used (Fig. 1C). The observed eIF4A dependency indicated that CR-31-B (-) may have antiviral activity against the newly emerging SARS-CoV-2 in cell culture.

**Figure 1:**
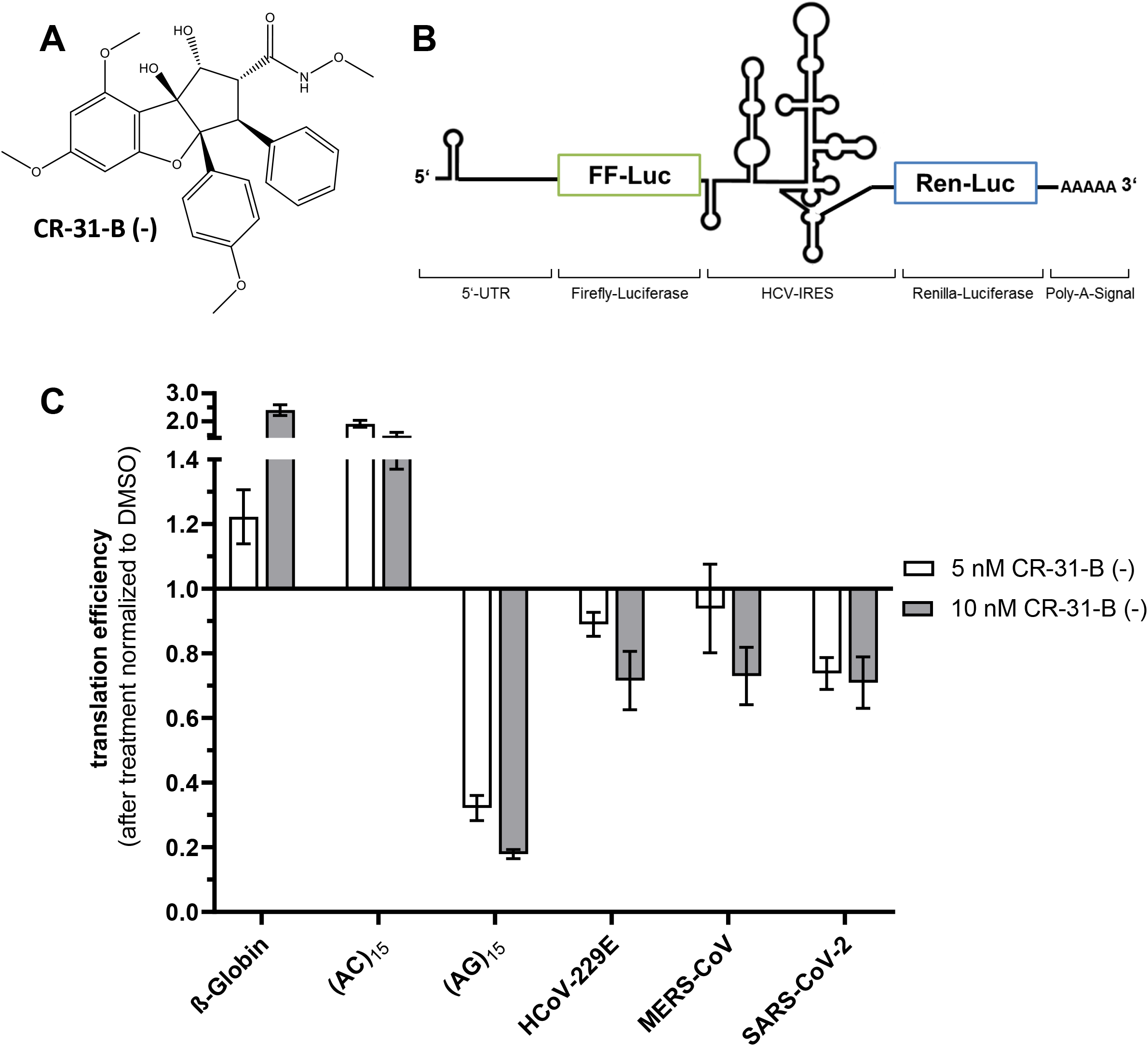
Effect of CR-31-B (-) on reporter gene expression from constructs containing different 5’-UTRs. **(A)** Structure of the synthetic rocaglate CR-31-B (-). **(B)** Schematic illustration of the dual luciferase reporter construct used to determine eIF4A-dependent translation of coronavirus 5’-UTRs. **(C)** Effects of 5 and 10 nM CR-31-B (-) on reporter gene expression in the context of 5’-UTRs from three human coronaviruses, HCoV-229E, MERS-CoV and SARS-CoV-2. The 5’-UTR of the human β-globin mRNA and the unstructured (AC)_15_ sequence served as negative controls, while the (AG)_15_ polypurine sequence served as a positive control. Experiments were performed with at least three independent biological replicates.

### 3.2. In vitro antiviral effect of CR-31-B (-) against SARS-CoV-2 in African green monkey Vero E6 cells

To analyze possible antiviral effects of CR-31-B (-) in cell culture, African green monkey Vero E6 cells were used because SARS-CoV-2 has previously been shown to grow to high titers in this cell line (Ogando et al., 2020). Cytotoxicity of CR-31-B (-) was determined in an MTT assay after treating Vero E6 cells with increasing concentrations of CR-31-B (-) for 24 h (Fig. 2A). As shown in Fig. 2A, no major cytotoxicity was detected for concentrations of up to 100 nM, with cell viability being reduced by about 10 - 25 % at this highest concentration tested (Fig. 2A). To determine antiviral activity of CR-31-B (-) against SARS-CoV-2, Vero E6 cells were infected with this virus at an MOI of 0.1 pfu/cell for 1 h and afterwards incubated in medium containing the different concentrations of CR-31-B (-). At 24 h p.i., cell culture supernatants were collected and virus titers were determined by plaque assay. The production of infectious SARS-CoV-2 progeny was found to be reduced in a dose-dependent manner with an EC_50_ concentration of ~1.8 nM (Fig. 2B), which is in a similar range with the CR-31-B (-) EC_50_ values reported previously for other coronaviruses (~2.9 nM for HCoV-229E; ~1.9 nM for MERS-CoV) (Müller et al., 2020). Next, we analyzed the effect of CR-31-B (-) on SARS-CoV-2 protein accumulation and the formation of viral replication/transcription complexes in Vero E6 cells. Viral nucleocapsid (N) protein levels were found to be drastically reduced in the presence of 100 nM CR-31-B (-), and moderately reduced at concentrations of 10 or 1 nM (Fig. 3A). As expected, viral protein accumulation was not affected in the presence of 100 nM of the (+)-enantiomer CR-31-B (+) nor was it affected in cells treated with solvent only.

**Figure 2:**
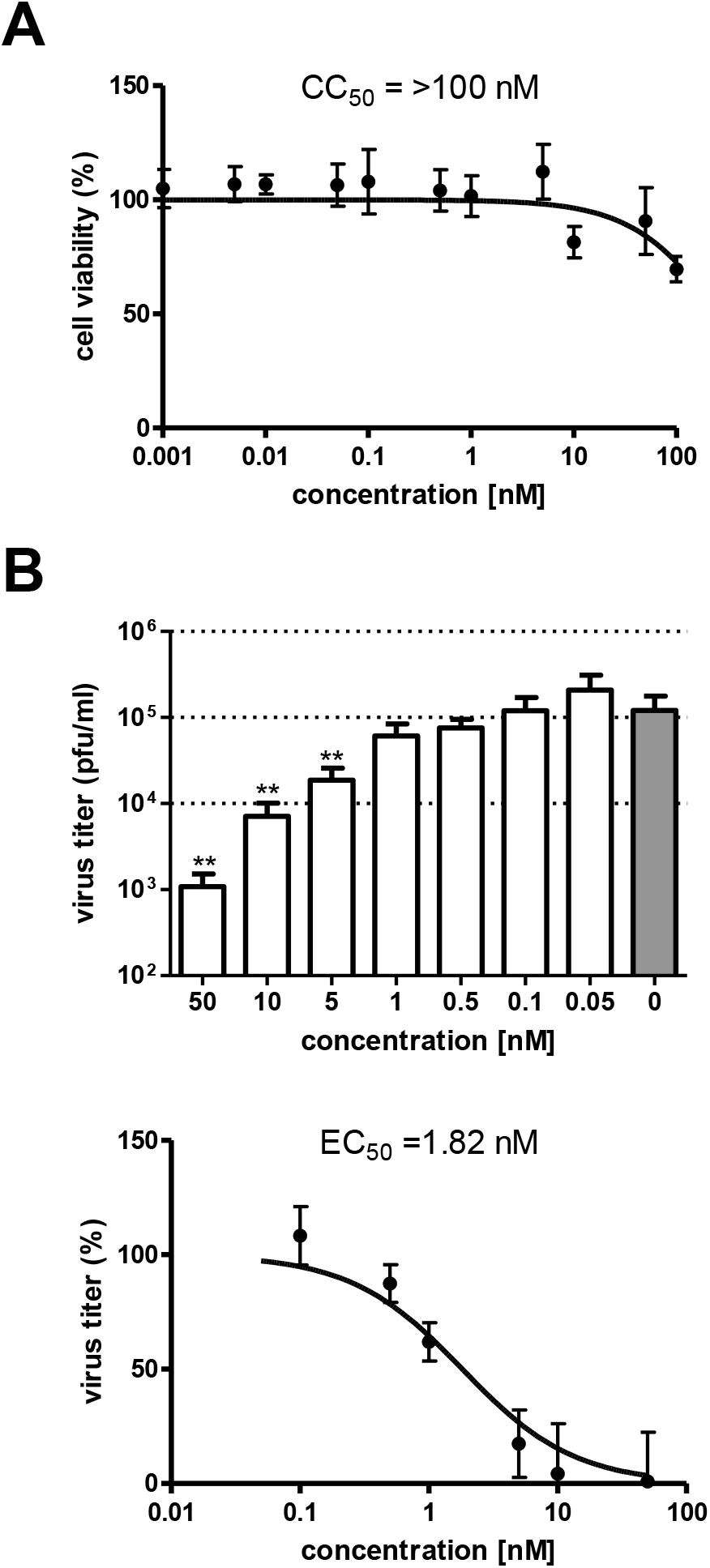
Dose-dependent antiviral activity of the synthetic rocaglate CR-31-B (-) in SARS-CoV-2 infected Vero E6 cells. **(A)** MTT assay of Vero E6 cells treated for 24 h with the indicated CR-31-B (-) concentrations. Cell viability (compared to that of untreated cells) was determined using the tetrazolium-based reagent. **(B)** SARS-CoV-2 titers in supernatants collected from infected Vero E6 cells (MOI = 0.1) treated with the indicated CR-31-B (-) concentrations were collected at 24 h p.i. (n=6) and virus titers were determined by plaque assay. Significance levels compared to the results for untreated cells were determined by the two-tailed Mann Whitney U-test and are indicated as follows: *, *P*<0.05; **, *P*<0.005. Data from six independent experiments were used to calculate the EC_50_ value by non-linear regression analysis.

In another set of experiments, we were able to show that the formation of SARS-CoV-2 replication/transcription complexes was impaired in infected cells treated with CR-31-B (-). As shown in Fig. 3B, immunofluorescence analysis using antibodies specific for double-stranded RNA (dsRNA), representing a viral RNA replication intermediate, revealed a profound reduction of replicative organelles active in viral RNA synthesis (Müller et al., 2018b).

**Figure 3:**
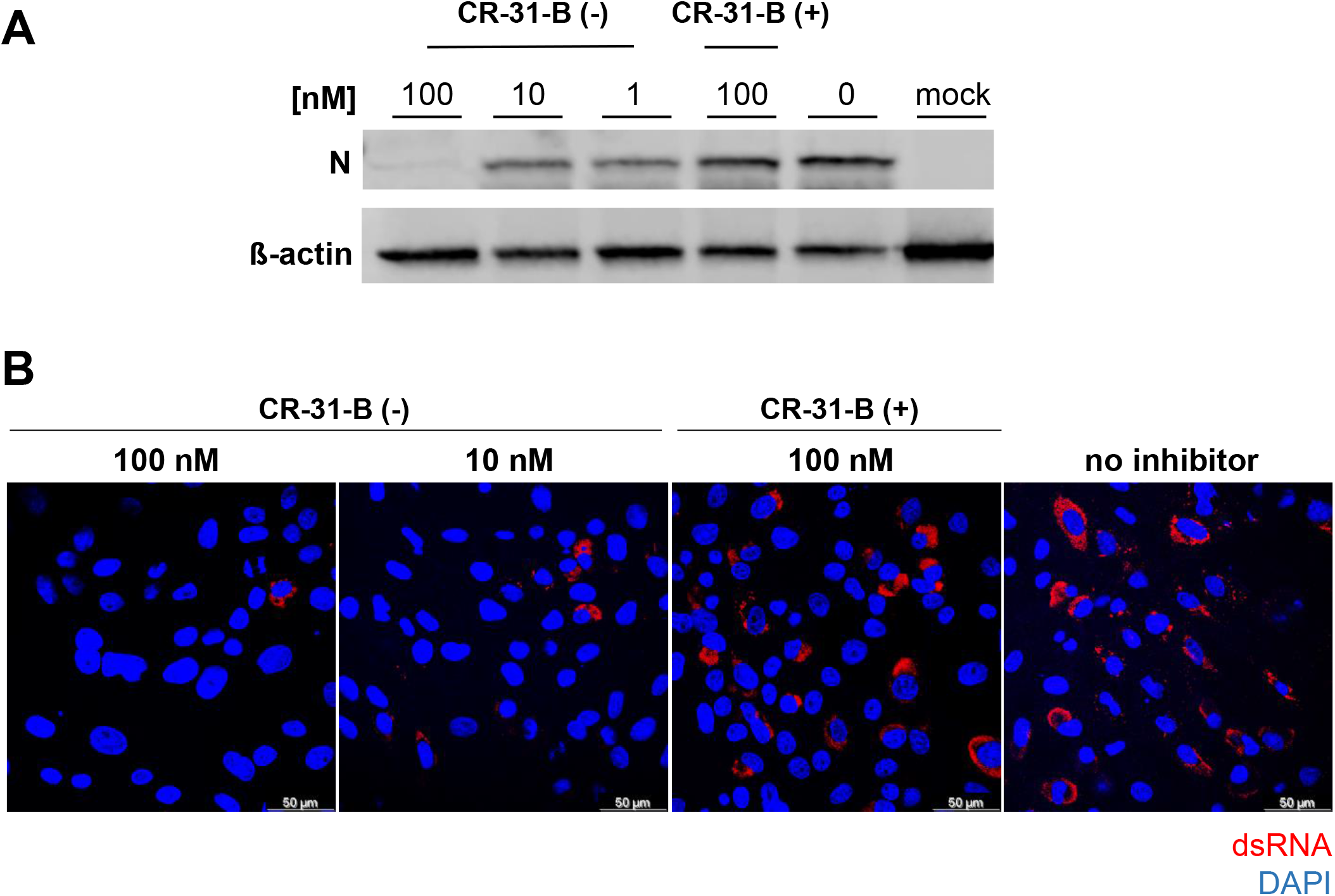
Antiviral activity of the synthetic rocaglate CR-31-B (-) in SARS-CoV-2-infected Vero E6 cells **(A)** Western blot analysis of SARS-CoV-2 N protein accumulation (top panel) in Vero E6 cells treated with the two enantiomers CR-31-B (-) and CR-31-B (+). β-actin (lower panel) was used as a loading control. **(B)** Immunofluorescence analysis to visualize the effects of CR-31-B (+) and CR-31-B (-) on viral dsRNA accumulation in SARS-CoV-2-infected Vero E6 cells. Cells were infected with an MOI of 1 and cultivated in medium containing the indicated CR-31-B concentrations. Cells were fixed at 12 h p.i. and analyzed by confocal microscopy using a dsRNA-specific antibody (red). DAPI staining of the DNA in the nucleus is shown in blue.

### 3.3. Antiviral activity of CR-31-B (-) and silvestrol against SARS-CoV-2 in an ex vivo human bronchial epithelial cell system

To further evaluate the antiviral potential of CR-31-B (-) in a biologically relevant *ex vivo* respiratory cell culture system, we analyzed air/liquid interface (ALI) cultures of differentiated primary normal human bronchial epithelial (NHBE) cells isolated from two different donors. ALI cultures are increasingly recognized as an excellent culture model mimicking the tracheobronchial region of the human respiratory tract and thus enabling respiratory infection research in a physiologically relevant cellular environment (Jonsdottir and Dijkman, 2016). Differentiated NHBE cells (Fig. 4A) were infected with SARS-CoV-2 (MOI = 3 pfu/cell) in the presence of inhibitor, CR-31-B (-) or silvestrol, the inactive enantiomer CR-31-B (+) or solvent control (untreated). Silvestrol was included as a reference in this experiment because this rocaglate has previously been tested extensively against a broad range of viruses. Silvestrol treatment reduced SARS-CoV-2 titers at different time points p.i. about 10 to 100-fold when used at a concentration of 10 nM, while virus replication in NHBE cells was completely abolished at a concentration of 100 nM (Fig. 4B). Next, we analyzed the effects of CR-31-B in NHBE cells. CR-31-B (-) reduced the production of infectious virus progeny by ~1.5 log steps at a concentration of 10 nM in differentiated NHBE cells obtained from two different donors. At 100 nM, CR-31-B (-) reduced SARS-CoV-2 titers to undetectable levels, whereas the inactive enantiomer CR-31-B (+) did not affect viral replication compared to the solvent control (Fig. 4C). Taken together, the data confirm a potent antiviral activity of CR-31-B (-) against this newly emerging coronavirus in a human *ex vivo* cell culture system. The compound proved to be active at nanomolar concentrations, similar to those reported previously for HCoV-229E (Müller et al., 2020).

**Figure 4:**
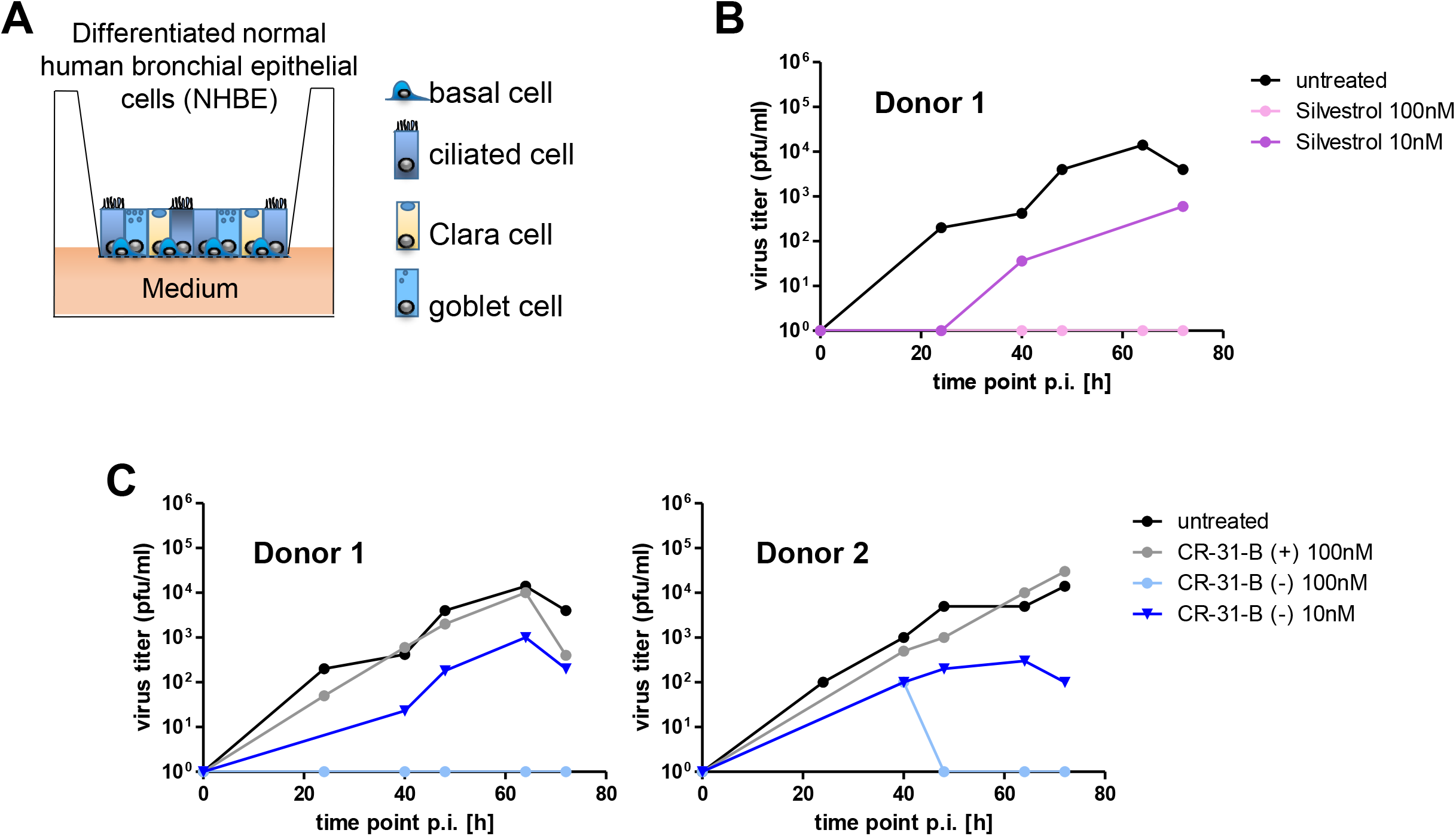
Comparison of antiviral effects of CR-31-B (-) and silvestrol using differentiated normal human bronchial epithelial (NHBE) cells infected with SARS-CoV-2. **(A)** NHBE cells were cultivated and differentiated into different cell types (Clara, ciliated, goblet and basal cells) under air-liquid interface conditions. **(B and C)** Differentiated NHBE cells were infected with SARS-CoV-2 (MOI = 3) and treated with **(B)** silvestroland **(C)** CR-31-B (-) or CR-31-B (+) at the indicated concentrations. SARS-CoV-2 titers in cell culture supernatants, collected at the indicated time points p.i. were determined by plaque assay.

## 4. Discussion

The ongoing SARS-CoV-2 pandemic, a coronavirus outbreak of zoonotic origin reminiscent of the SARS and MERS outbreaks from a few years ago, has led to an increased awareness of the need for first-line, broad-spectrum ‘pan-antivirals’ that can be used as stopgaps until vaccines and specific therapies become available to prevent or treat infections caused by newly emerging viruses.

We and others have focused on the discovery and development of antivirals that target host mechanisms critical to viral proliferation. The rationale behind this approach is twofold: first, host mechanisms are not virus-specific, but are used by a broad range of viruses, and second, targeting a host mechanism preempts the risk for developing resistance typically associated with targeting viral structures or mechanisms.

Several host-targeting approaches against SARS-CoV-2 are currently under investigation. Most of these represent efforts to repurpose approved drugs with known safety profiles. Promising candidates are for example inhibitors of SARS-CoV-2 entry (e.g. compounds that block the viral spike protein activating cellular transmembrane protease serine 2 (TMPRSS2), (for review see Xiu et al., 2020)), several immunomodulatory agents and also some kinase inhibitors that interfere with virus particle assembly (e.g. Janus kinase inhibitors) or with virus replication (imatinib). Angiotensin converting enzyme inhibitors or angiotensin receptor blockers, hydroxy methylglutaryl coenzyme A (HMG-CoA) reductase inhibitors and several other compounds have also been tested as single usage or in combination approaches (Lam et al., 2020, Al-Horani et al., 2020, Sharma et al., 2020, Santos et al., 2020), thus illustrating an increasing relevance of host-targeting antiviral therapeutic strategies to combat COVID-19.

Here, we have further expanded the antiviral toolbox against SARS-CoV-2 by characterizing the antiviral activity of the synthetic rocaglate CR-31-B (-), a specific inhibitor of eIF4A-dependent mRNA translation. We identified CR-31-B (-) as a potent and non-cytotoxic inhibitor of SARS-CoV-2 replication *in vitro* and *ex vivo*. The observed antiviral activities are directly comparable to those we have reported previously for other coronaviruses, namely HCoV-229E and MERS-CoV, as well as a range of highly pathogenic positive- and negative-sense single-stranded RNA viruses. This is to our knowledge the first report of a rocaglate inhibiting SARS-CoV-2 proliferation in a relevant *ex vivo* human bronchial cell system.

Against the backdrop of a number of reports we and others have published over the past three years documenting the broad-spectrum antiviral activity of CR-31-B (-) and related rocaglates, the data we report here strengthen the case for the potential use of rocaglates as first-line, broad-spectrum ‘pan-antivirals’. The necessary next step is the *in vivo* evaluation of CR-31-B (-) and other rocaglates to confirm safety and determine appropriate dosing regimens. Extensive preclinical work with CR-31-B (-), zotatifin, and other rocaglates as cancer therapeutics is *a priori* highly encouraging for the potential antiviral application of rocaglates in humans. Minimal or no toxicities seen following long-term dosing regimens in animal models, favorable ADME- and bioavailability profiles (Wolfe et al., 2014, Chan et al., 2019, Rodrigo et al., 2012), and the advancement of at least one rocaglate into the clinic for cancer, zotatifin, support the potential for this class of molecules to be used as antivirals in humans. At this point it is too early to speculate whether rocaglates could be applied prophylactically or therapeutically, but ongoing *in vivo* studies with CR-31-B (-) in relevant animal models of virus infection will provide answers to these questions.

In summary, the path to clinical implementation is still long and will most likely require further optimization of CR-31-B (-) or other lead rocaglates currently under investigation. Nevertheless, the consistent antiviral efficacy and low toxicity of this class of compounds across viruses both *in vitro* and *ex vivo* and the more advanced parallel efforts to harness their activity as cancer therapeutics suggest that rocaglates represent a potentially powerful addition to the toolbox of interventions available to public health authorities to counter new and emerging RNA virus outbreaks globally.

## 5. Acknowledgements

We like to thank Ulrike Wend for technical assistance.

